# TgIF2K-B is an eIF2α kinase in *Toxoplasma gondii* that responds to oxidative stress and optimizes pathogenicity

**DOI:** 10.1101/2020.09.24.312744

**Authors:** Leonardo Augusto, Jennifer Martynowicz, Parth H. Amin, Kenneth R. Carlson, Ronald C. Wek, William J. Sullivan

## Abstract

*Toxoplasma gondii* is an obligate intracellular parasite that persists in its vertebrate hosts in the form of dormant tissue cysts, which facilitate transmission through predation. The parasite must strike a balance that allows it to disseminate throughout its host without killing it, which requires the ability to properly counter host cell defenses. For example, oxidative stress encountered by *Toxoplasma* is suggested to impair parasite replication and dissemination. However, the strategies by which *Toxoplasma* mitigates oxidative stress are not completely clear. Among eukaryotes, environmental stresses induce the integrated stress response via phosphorylation of a translation initiation factor, eIF2. Here, we show that the *Toxoplasma* eIF2 kinase, TgIF2K-B, is activated in response to oxidative stress and affords protection. Knockout of TgIF2K-B, Δ*tgif2k-b*, disrupted parasite responses to oxidative stresses and enhanced replication, diminishing the ability of the parasite to differentiate into tissue cysts. In addition, parasites lacking TgIF2K-B exhibited resistance to activated macrophages and showed greater virulence in an *in vivo* model of infection. Our results establish that TgIF2K-B is essential for *Toxoplasma* responses to oxidative stress, which are important for the parasite’s ability to establish persistent infection in its host.

**IMPORTANCE:** *Toxoplasma gondii* is a single-celled parasite that infects nucleated cells of warm-blooded vertebrates, including one-third of the human population. The parasites are not cleared by the immune response and persist in the host by converting into a latent tissue cyst form. Development of tissue cysts can be triggered by cellular stresses, which activate a family of TgIF2 kinases to phosphorylate the eukaryotic translation initiation factor, TgIF2α. Here, we establish that the TgIF2 kinase TgIF2K-B is activated by oxidative stress and is critical for maintaining oxidative balance in the parasite. Depletion of TgIF2K-B alters gene expression, leading to accelerated growth and a diminished ability to convert into tissue cysts. This study establishes that TgIF2K-B is essential for the parasite’s oxidative stress response and its ability to persist in the host as a latent infection.

## Introduction

*Toxoplasma gondii* is a protozoan parasite that infects nucleated cells of warm-blooded vertebrates, including up to one-third of the human population (1). *Toxoplasma* is an obligate intracellular pathogen that replicates inside of a parasitophorous vacuole (tachyzoites) that forms an interface between the parasites and their host cell (2). One of the features that facilitated *Toxoplasma*’s omnipresence in the animal kingdom is the ability to persist in its host as latent tissue cysts (bradyzoites) that serve to disseminate the parasite to new hosts via predation (3). *Toxoplasma* has thus evolved mechanisms to balance parasite metabolism, replication, and cyst formation to allow rapid dissemination and encystment throughout host tissues without killing the host (4).

In addition to facilitating transmission, tissue cysts afford *Toxoplasma* protection from immune defenses, thus preventing the parasite from being eliminated from the host. However, a consequence of *Toxoplasma* persistence as a cyst is diminished parasite replication. Rapidly proliferating tachyzoites respond to environmental stresses by slowing parasite growth and by triggering differentiation into bradyzoites (3). A critical inverse relationship between the rate of cell proliferation and resistance to environmental stresses has been noted among other unicellular eukaryotes (5). This balance provides *Toxoplasma* with resistance to physiological and environmental stresses, and ensures optimal infection and transmission into new hosts.

An important mechanism for mitigating stress damage in eukaryotes is the integrated stress response (ISR). The ISR features a family of related protein kinases that phosphorylate the α subunit of eukaryotic initiation factor 2 (eIF2α) in response to a variety of stresses. Phosphorylation of eIF2α dampens global translation, which conserves nutrients and energy, while promoting preferential expression of genes involved in stress remediation (6). Phosphorylation of *Toxoplasma* eIF2α (TgIF2α-P) occurs in response to environmental stresses and is a contributing factor in the formation of tissue cysts (7, 8). Four TgIF2α kinases have been identified in the parasite, designated A through D (9). TgIF2K-A is localized to the parasite endoplasmic reticulum (ER) and responds to perturbations in this organelle, processes analogous to the eIF2α kinase PERK (EIFAK3/PEK) (10). TgIF2K-C and -D are related the eIF2α kinase GCN2 (EIFAK4) and respond to amino acid starvation and stresses of the extracellular environment, respectively (11–13). The function of TgIF2K-B remains unknown.

In this study, we addressed the functions of TgIF2K-B in *Toxoplasma* replication, stress responses, and pathogenesis. As *Toxoplasma* is an aerobic parasite that needs to limit molecular damage caused by generation of excessive reactive oxygen species (ROS), including those generated by host defense mechanisms (14), we hypothesized that TgIF2K-B functions in antioxidation responses. We generated a genetic knockout for use in *in vitro* and *in vivo* host model systems to show that TgIF2K-B is essential for the activation of catalase antioxidative responses, which affects tachyzoite replication and bradyzoite conversion. Our results suggest that control of antioxidant responses is central to the parasite’s growth rate, which is a critical determinant in the ability of the parasite to persist in its host in a latent form. We conclude that TgIF2K-B is a novel sensor protein governing the rate of *Toxoplasma* replication and cyst formation through translational control mechanisms involving TgIF2α phosphorylation.

## Results

### Knockout of TgIF2K-B accelerates *Toxoplasma* replication

We have previously shown that TgIF2K-B is localized to the parasite cytosol and can phosphorylate TgIF2α *in vitro* (8). To address stress activation of TgIF2K-B and the role of this eIF2α kinase in parasite physiology and differentiation, we generated a knockout parasite in type II ME49 strain using CRISPR/Cas9 methods (15). The knockout was created through genetic insertion of a modified allele of DHFR that confers resistance to pyrimethamine into the first exon of the TgIF2K-B locus (16) **(Fig. 1A)**. PCR primers upstream of the insertion cassette (P1) were used to amplify the genomic sequences and primers inside of DHFR and in the TgIF2K-B gene were used to amplify across the DHFR insertion (P2). The PCR products indicated the anticipated DHFR insertion, generating ME49 Δ*tgif2k-b* parasites **(Fig. 1B)**. In addition, we confirmed the loss of *TgIF2K-B* mRNA and protein in the Δ*tgif2k-b* by RT-qPCR **(Fig. 1C)** and by immunoblot using a specific antibody (8) **(Fig. 1D)**.

**Figure 1.**
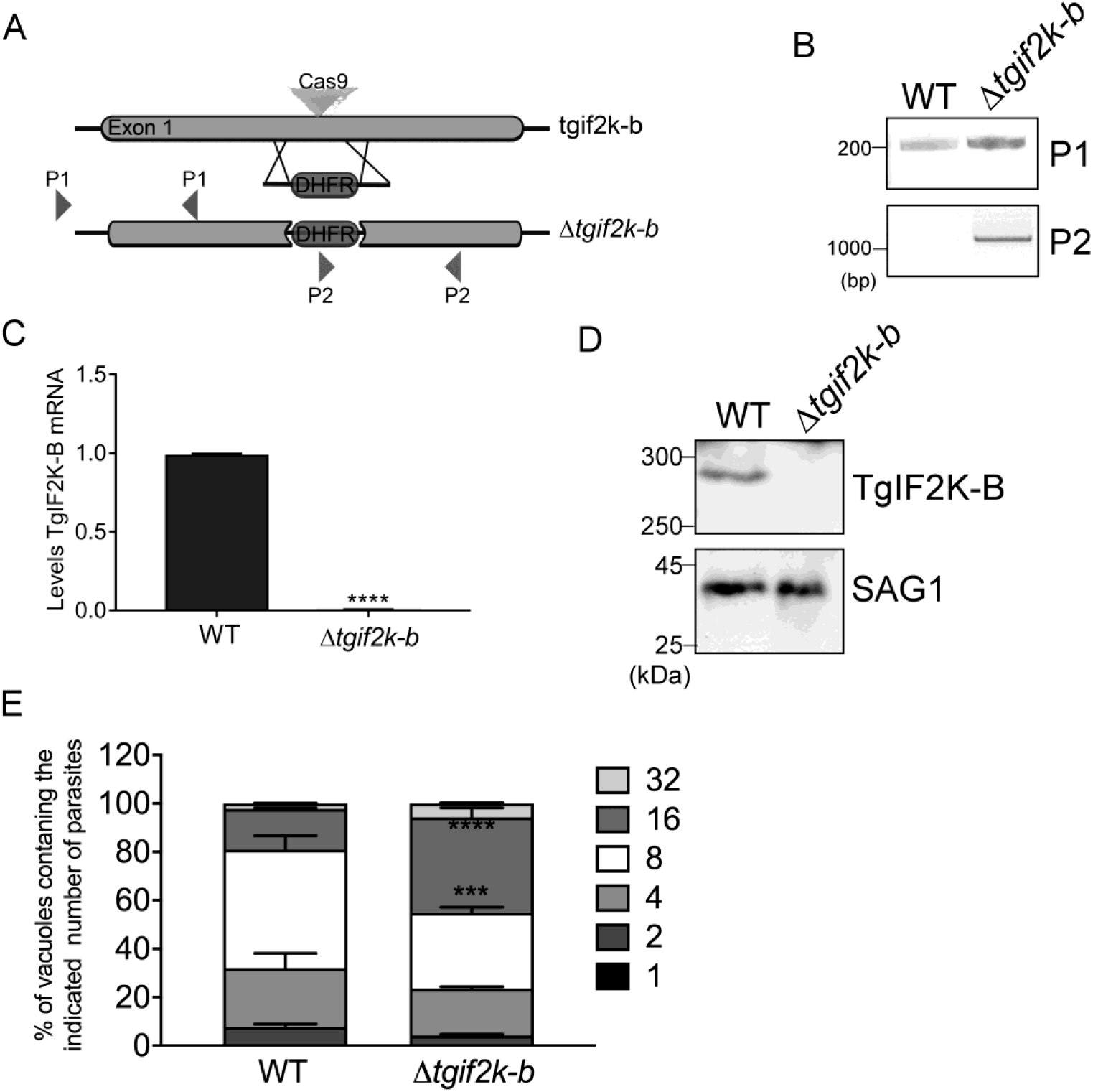
Disruption of *TgIF2K-B* gene in ME49 strain accelerates replication. **(A)** Strategy used to the disrupt the *TgIF2K-B* gene involved using CRISPR/Cas9 to insert a modified DHFR allele that encodes pyrimethamine resistance into exon 1. The indicated primers pairs, P1 and P2, were used to confirm the disrupted locus in drug-resistant clones. **(B)** Genomic DNA isolated from WT and Δ*tgif2k-b* parasites was used to assess structure of *TgIF2K-B* locus by PCR with P1 and P2 primer pairs. PCR products were separated by agarose gel electrophoresis and the size of bands are indicated in base pairs (bp). **(C)** Levels of *TgIF2K-B* mRNA were measured by RT-qPCR for WT and Δ*tgif2k-b* parasites (±SD, n=3). ****p<0.0001. **(D)** Immunoblot of WT and Δ*tgif2k-b* lysates for TgIF2K-B protein. SAG1 was probed as a loading control. Molecular weights are indicated in kDa. **(E)** At 36 hours post-infection, the number of parasites in 250 random vacuoles was counted and presented as a percentage of the total number of vacuoles examined (±SD, n=3). ***p<0.005 and ****p<0.0001.

It was noted that parasite clones lacking TgIF2K-B appeared to grow faster than wild-type (WT), which was confirmed by counting the number of parasites per vacuole in infected human foreskin fibroblast (HFF) cells. Δ*tgif2k-b* parasites produced significantly more vacuoles containing 16 and 32 parasites at the 36 hour time point than WT (Fig. 1E). These results suggest that the TgIF2K-B pathway plays a role in the regulation of parasite replication.

### TgIF2K-B participates in the oxidative stress response

To address the function of TgIF2K-B during infection and assess which stress condition(s) activates TgIF2K-B, we treated extracellular WT and Δ*tgif2k-b* parasites with the following stress agents: thapsigargin (TG), an ER stress inducer that leads to activation of TgIF2K-A (10), halofuginone (HF), which inhibits the aminoacylation of tRNA^Pro^ and is also a potent inducer of the eIF2α kinase GCN2, and sodium arsenite (Ars), which triggers general oxidative stress. We detected increased levels of TgIF2α-P in WT parasites exposed to each of the three different stress conditions **(Fig. 2A)**. By contrast, Δ*tgif2k-b* parasites displayed minimal levels of TgIF2α-P upon treatment with arsenite (Ars), whereas the other stress conditions showed robust TgIF2α-P **(Fig. 2A)**. As expected after phosphorylation of TgIF2α, global protein synthesis was lowered by arsenite stress in WT parasites; however, it remained unaffected in Δ*tgif2k-b* parasites exposed to arsenite **(Fig. 2B and Fig. S1)**. Also of note is the higher level of basal protein synthesis in Δ*tgif2k-b* parasites compared to WT, consistent with the demands of faster replication **(Fig. 2B)**. In support of the idea that oxidative stress induced by arsenite is critical for induction of TgIF2α-P, co-treatment of the parasites with arsenite and the antioxidant N-acetyl cysteine (Nac) led to a sharp decrease in TgIF2α-P levels in the WT parasites **(Fig. 2C)**. We conclude that TgIF2K-B is an eIF2α kinase activated by oxidative stress in *Toxoplasma*.

**Figure 2.**
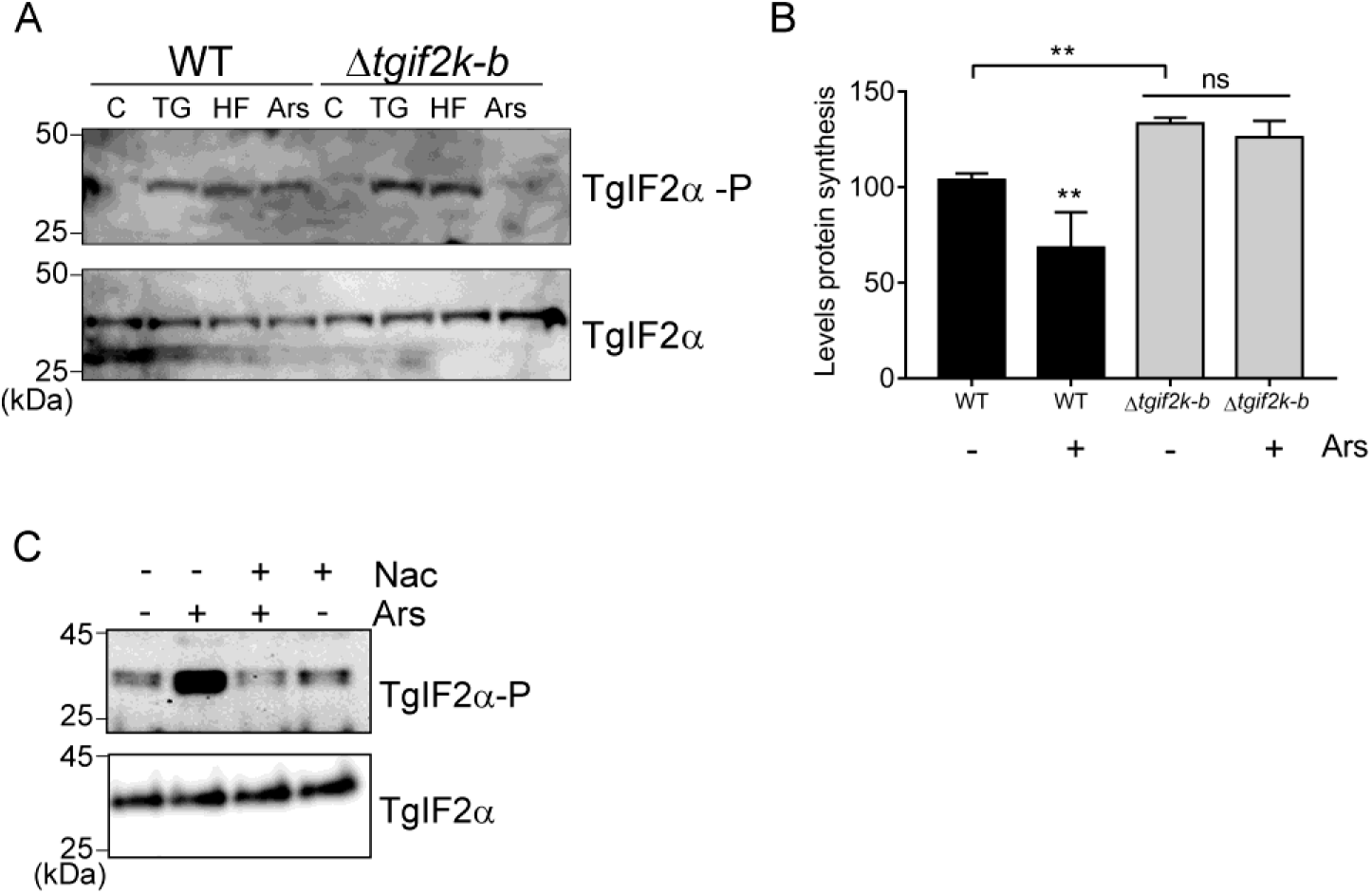
TgIF2K-B responds to oxidative stress. **(A)** Extracellular WT and Δ*tgif2k-b* tachyzoites were treated with 1 μM thapsigargin (TG), 50 nM halofuginone (HF), or 5 μM sodium arsenite (Ars) for 2 hours at 37°C. Parasites were harvested and the levels of phosphorylated TgIF2α (TgIF2α-P) and total TgIF2α were measured by immunoblot. **(B)** WT and *Δtgif2k-b* tachyzoites were subjected to 5 μM Ars or vehicle for 2 hours, then treated with puromycin for 15 min. Lysates were then analyzed by immunoblot with puromycin-specific antibody. Levels of translation are represented as a bar graph with amounts shown relative to WT without Ars treatment (±SD, n=3). **p<0.01, ns = not significant. **(C)** WT parasites were treated with 5 μM Ars for 2 hours in the presence or absence of 20 μM N-acetylcysteine (Nac), then TgIF2α-P and total TgIF2α were assessed by immunoblot.

To address the transcriptome changes resulting from TgIF2K-B deletion, we performed RNA-seq analyses of WT and Δ*tgif2k-b* parasites. Differentially expressed genes in Δ*tgif2k-b* parasites were selected using log2fold change of ≤ or ≥ ±1 and p-adjusted values of ≤ 0.05 as detailed in Methods. Loss of TgIF2K-B resulted in induction of 1,162 genes and lowered expression of 1,686 genes relative to WT parasites **(Fig. 3A-B)**. The RNA-seq analyses showed loss of TgIF2K-B led to enhanced levels of *Toxoplasma* superoxide dismutase-1 (Tg*SOD1*) mRNA and downregulation of genes related to antioxidation, including PRX2, thioredoxin, and oxidoreductases **(Fig. 3A)**. These results suggest that TgIF2K-B directs expression of genes involved in mitigating oxidative damage.

**Figure 3.**
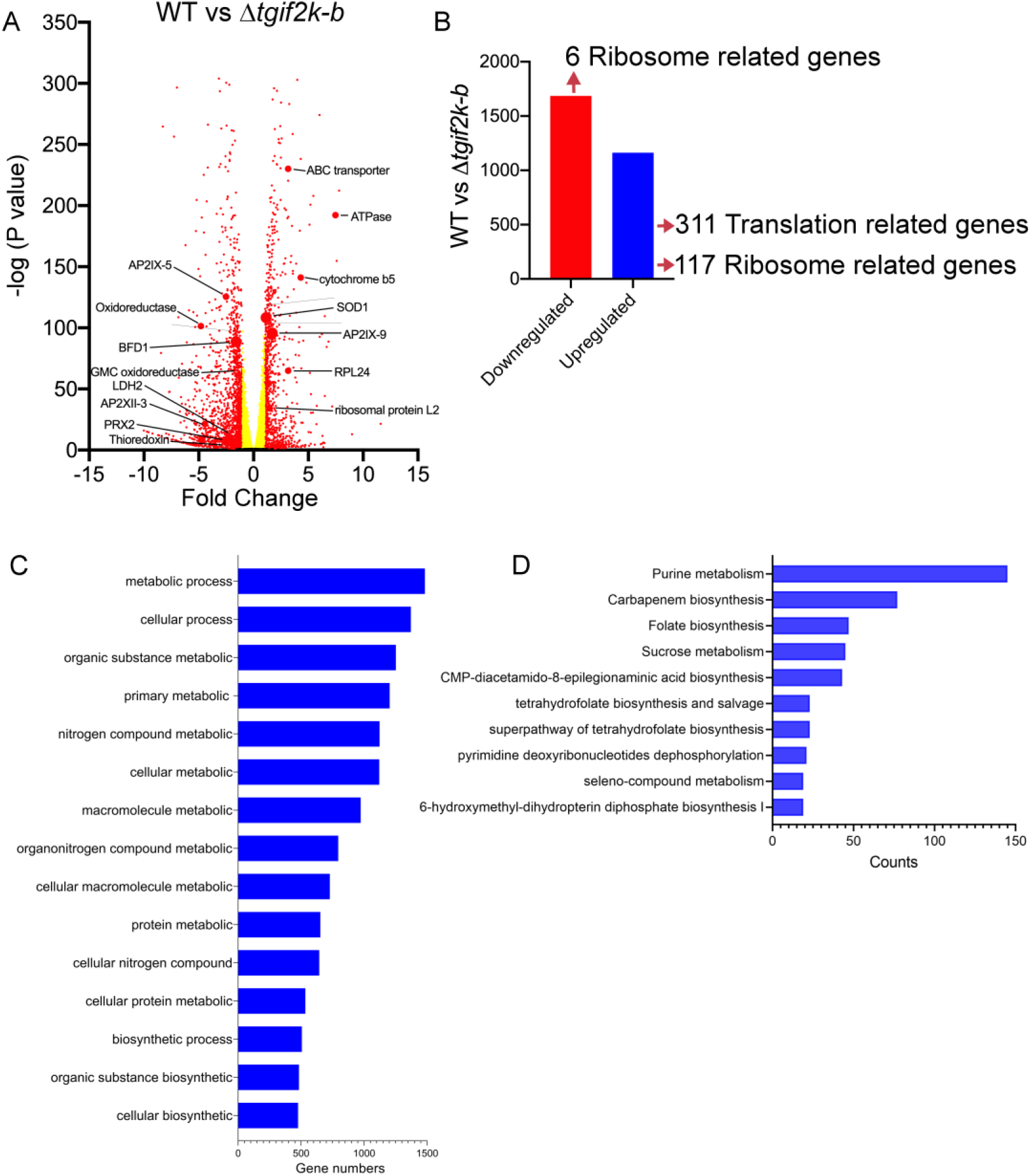
RNA-seq analysis of WT and *Δtgif2k-b* parasites. RNA-seq and differential expression of genes in *Δtgif2k-b* parasites compared to WT parasites. **(A)** Volcano plots showing transcription profile of *Δtgif2k-b* parasites compared to WT parasites (WT vs *Δtgif2k-b*). **(B)** Number of up- or down-regulated genes in WT vs *Δtgif2k-b*; 1,162 genes were identified as significantly upregulated (blue) and 1,686 genes as significantly downregulated (red) in *Δtgif2k-b* parasites. The analyses were based on biological replicates using fold change of 1.0 and p-value ≤ 0.05. **(C)** Gene ontology enrichment analysis for biological processes (using ToxoDB) of upregulated genes in *Δtgif2k-b* compared to WT parasites. **(D)** Gene ontology enrichment analysis for metabolic pathways (using ToxoDB) of upregulated genes in *Δtgif2k-b* parasites. In panels C and D, the length of the bars indicates the number of genes identified for the indicated processes and pathways.

It is noteworthy that loss of TgIF2K-B resulted in upregulation of 311 genes involved in protein synthesis, including 117 ribosome-related genes, whereas 6 ribosome-related genes were downregulated **(Fig. 3B)**. These results further support that there are higher levels of protein synthesis in Δ*tgif2k-b* parasites that would contribute to increased replication **(Fig. 1E and 2B)**. Furthermore, the absence of TgIF2K-B resulted in enhanced expression of more than 1,000 genes involved in metabolic processes, including those involved in purine and carbohydrate metabolism and folate biosynthesis **(Fig. 4C-D, Fig. S2 and S3)**.

**Figure 4.**
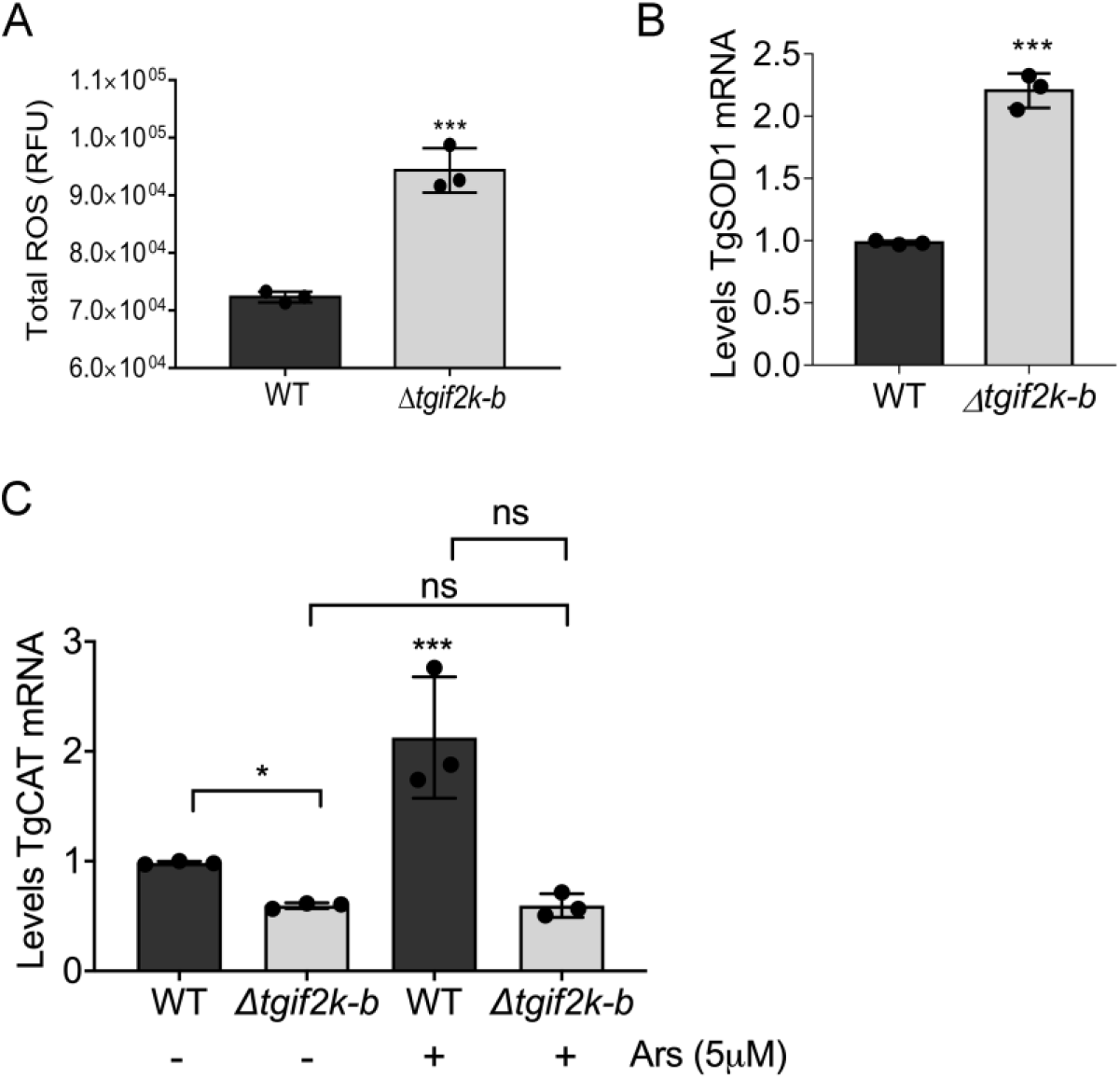
Loss of TgIF2K-B disrupts the antioxidant response. **(A)** Reactive species (ROS) were measured in WT and Δ*tgif2k-b* parasites using OxiSelect ROS/RNS; values were normalized to total protein (±SD, n=3). ***p<0.005. **(B)** *TgSOD1* mRNA levels were measured by RT-qPCR in WT and Δ*tgif2k-b* parasites (±SD, n=3). ***p<0.005. **(C)** *TgCAT*mRNA levels were measured by RT-qPCR in WT and Δ*tgif2k-b* parasites subjected to 5 μM Ars or vehicle for 2 h at 37°C (±SD, n=3). * p<0.05, ***p<0.005, ns = not significant.

To more fully address the redox functions of TgIF2K-B, we measured ROS levels in WT and knockout parasites. Results indicate that deletion of TgIF2K-B leads to significantly higher levels of ROS compared to WT parasites **(Fig. 4A)**. It has been previously suggested that ROS act as second messengers regulating the balance between cell proliferation and cell cycle (17, 18). An early step in antioxidation responses involves SOD, which generates H_2_O_2_ that will subsequently be decomposed by catalase into oxygen and water. We first measured Tg*SOD1* mRNA levels by RT-qPCR, confirming that they are enhanced in Δ*tgif2k-b* parasites as detected in the RNA-seq **(Fig. 3A, 4B)**. We next measured expression of catalase (*TgCAT*) mRNA in WT and Δ*tgif2k-b* parasites after treatment with sodium arsenite for 2 hours. While WT parasites increase Tg*CAT* mRNA levels upon arsenite treatment, Δ*tgif2k-b* parasites fail to do so **(Fig. 4C)**. These results support that TgIF2K-B activation and phosphorylation of TgIF2α affects expression of antioxidant genes.

To address the transcriptome changes upon arsenite treatment and determine the contributions of TgIF2K-B on this gene expression network, we treated extracellular WT and Δ*tgif2k-b* tachyzoites for 2h with 5 μM arsenite and then performed RNA-seq analyses. The analyses indicated that the expression of 780 genes were enhanced in WT parasites in response to arsenite treatment and 452 genes were downregulated **(Fig. 5A, B and C)**. A major cell process enhanced in WT parasites in response to arsenite involves oxidation-reduction **(Fig. 5D)**, including induction of catalase as noted earlier **(Fig. 4C)**. By contrast, a majority of downregulated genes by arsenite treatment in WT parasites are involved metabolism, including genes related to acetyl-coA carboxylase, glycosyl transferase, beta-ketoacyl-acyl carrier protein synthase, beta-ketoacyl-acyl carrier protein synthase, and amino acid transporters **(Fig. 5E and Table S4)**. We also compared the RNA-seq analyses between WT and Δ*tgif2k-b* parasites in response to arsenite treatment. In sharp contrast to the hundreds of genes altered in the WT response, Δ*tgif2k-b* parasites only upregulated 15 genes and downregulated 16 genes **(Fig. 5B and C)**. WT and Δ*tgif2k-b* parasites shared only 2 upregulated genes in response to arsenite **(Fig. 5B).** The results emphasize that the loss of TgIF2K-B greatly dysregulates the parasite’s response to oxidative stress.

**Figure 5.**
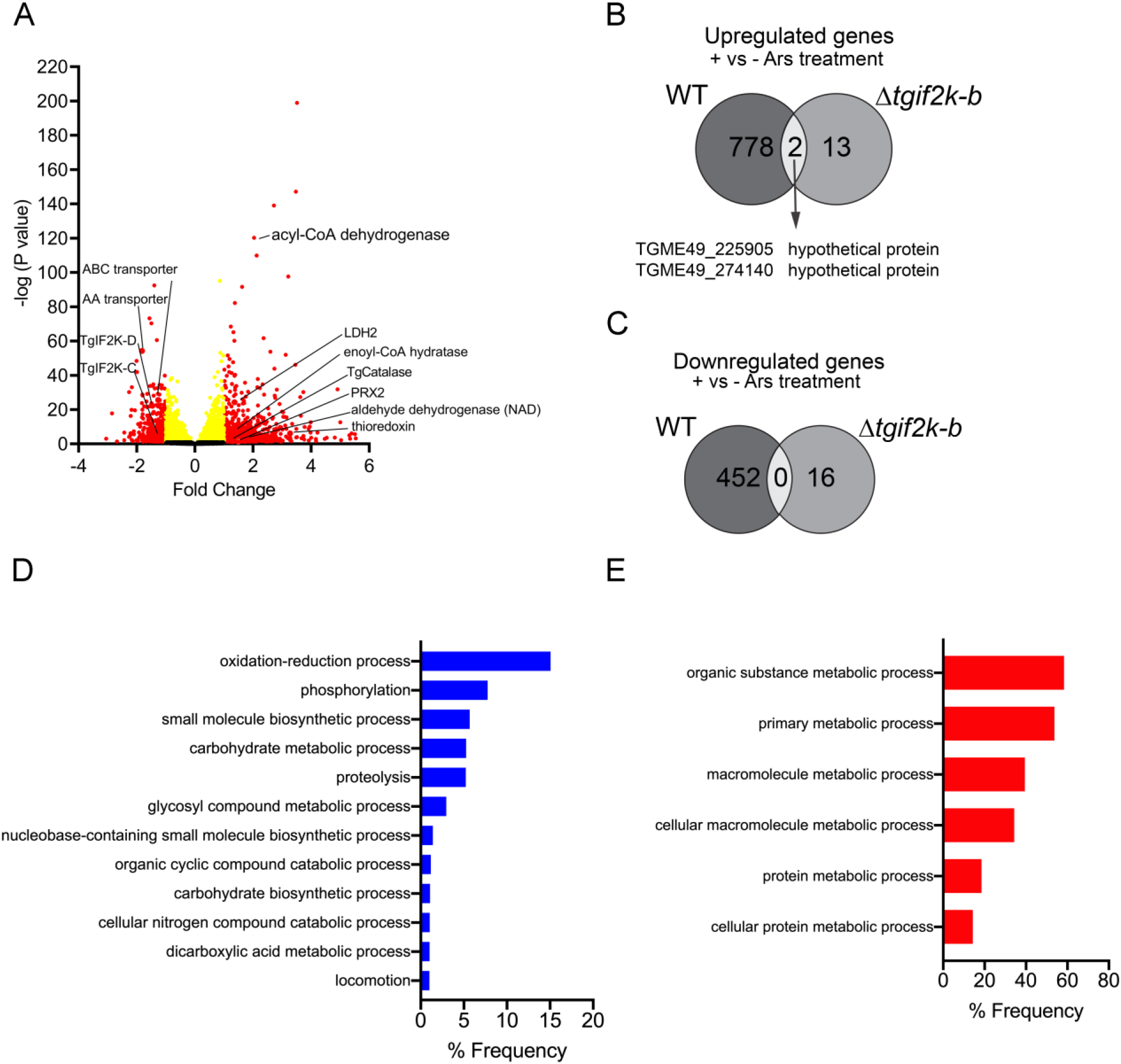
RNA-seq analysis of WT and *Δtgif2k-b* parasites treated with sodium arsenite. **(A)** Volcano plots showing transcription profile of WT parasites treated with Ars for 2 hours compared to WT parasites untreated. **(B-C)** WT and *Δtgif2k-b* parasites were treated with arsenite or vehicle for 2 hours and processed for RNA-seq. Numbers of differentially expressed genes were compared between WT and *Δtgif2k-b* parasites +/- arsenite. Gene Ontology enrichment analysis was used to identify cellular processes that were **(D)** upregulated or **(E)** downregulated in WT parasites +/- arsenite.

### Loss of TgIF2K-B leads to parasite resistance during macrophage infection

We next investigated how the compromised ability to respond to oxidative stress might affect parasites in macrophages, which produce ROS as a defense mechanism against *Toxoplasma* (19, 20). We addressed this by analyzing replication of WT and Δ*tgif2k-b* parasites in activated macrophages. Prior to infection, we stimulated J774.1 macrophages with LPS for 24 h; we then counted the number of intracellular parasites 4 days post-infection using the PCR-based B1 assay (21). While the number of WT parasites was significantly decreased in activated macrophages, replication of Δ*tgif2k-b* parasites was unimpeded **(Fig. 6A)**. These results suggest that TgIF2K-B is important for sensing oxidative stress generated by host immune cells, and the inability to do so allows tachyzoite replication to proceed unfettered.

**Figure 6.**
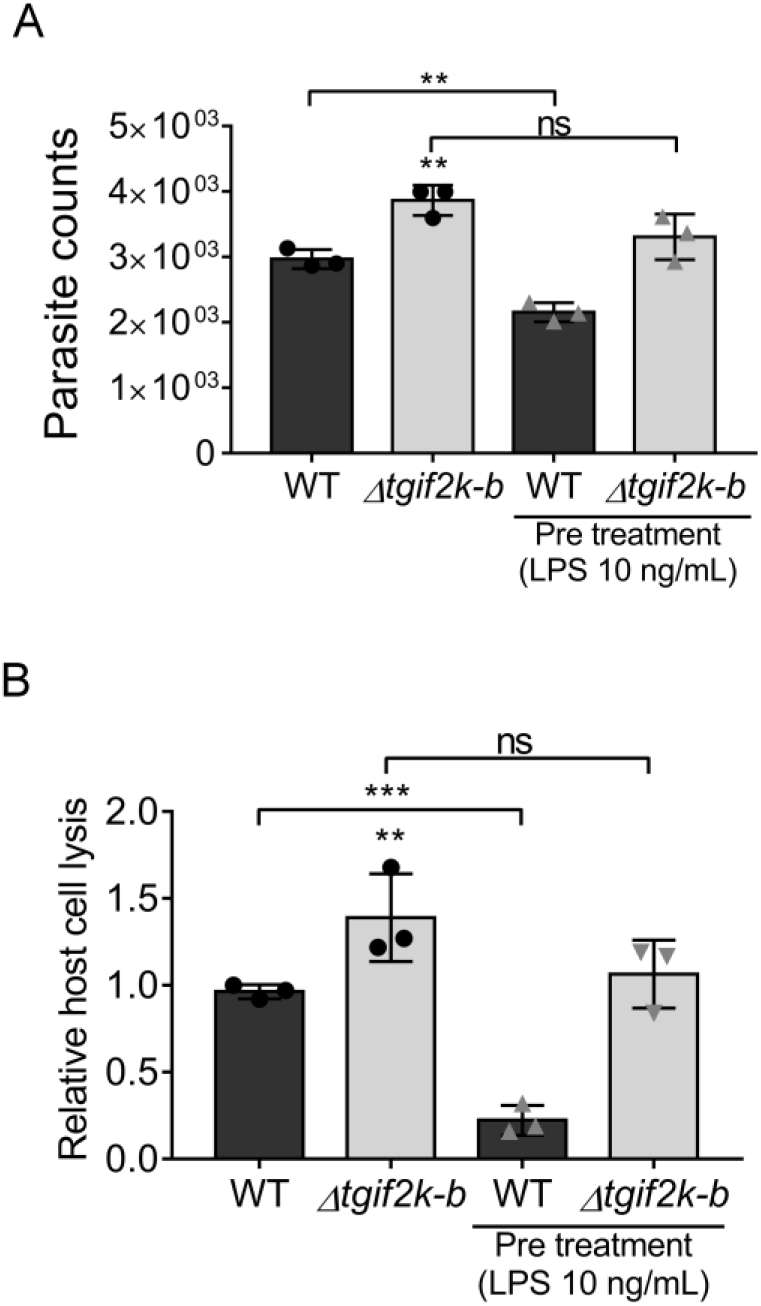
TgIF2K-B governs parasite replication in macrophages. **(A)** Tachyzoites were allowed to infect J774.1 macrophages that were activated by LPS for 24 hours. Four days post-infection, parasite replication was measured using the PCR-based B1 assay (±SD, n=3). **p<0.01, ns = not significant. **(B)** Tachyzoites were cycled through activated macrophages for 2 days before being harvested by syringe lysis to then inoculate HFF monolayers. After 6 days, the degree of HFF host cell lysis was quantified (±SD, n=3). **p<0.01, ***p<0.001.

In addition, we measured parasite viability following infection of macrophages that had been activated or not. Mock or LPS-stimulated J774.1 macrophages were infected with WT or Δ*tgif2k-b* parasites for 2 days, at which point parasites were harvested and used to infect an HFF cell monolayer **(Fig. 6B)**. After 6 days, the area of parasite plaques in the HFF monolayer was measured for each condition. Results show that cycling through macrophages significantly reduced HFF infectivity of WT tachyzoites, but the Δ*tgif2k-b* parasites displayed resistance to this insult **(Fig. 6B)**. The effect was more pronounced when the parasites were cycled through activated macrophages. These findings further support the idea that TgIF2K-B senses oxidative stresses and triggers a response that slows parasite growth to manage the stress.

### TgIF2K-B is a sensor protein governing replication rate and cyst formation

As phosphorylation of TgIF2α accompanies bradyzoite differentiation and tissue cyst formation (8), we hypothesized that parasites lacking TgIF2K-B may show deficits in tissue cyst formation. WT and Δ*tgif2k-b* parasites were allowed to spontaneously differentiate in HFF cells. After 5 days, Δ*tgif2k-b* cultures contained <10% tissue cysts whereas WT cultures contained ~50% **(Fig. 7A)**. In addition, the tissue cysts generated by the Δ*tgif2k-b* parasites were twice the size of those produced by WT parasites **(Fig. 7B and C)**. We propose that the increased replication rate of Δ*tgif2k-b* tachyzoites hinders bradyzoite conversion; while cyst wall proteins are secreted in sufficient quantity to interact with *Dolichos* lectin, the parasites inside continue dividing, resulting in larger cysts than WT. These results suggest that TgIF2K-B is a critical sensor that balances developmental transitions between replication and latency.

**Figure 7.**
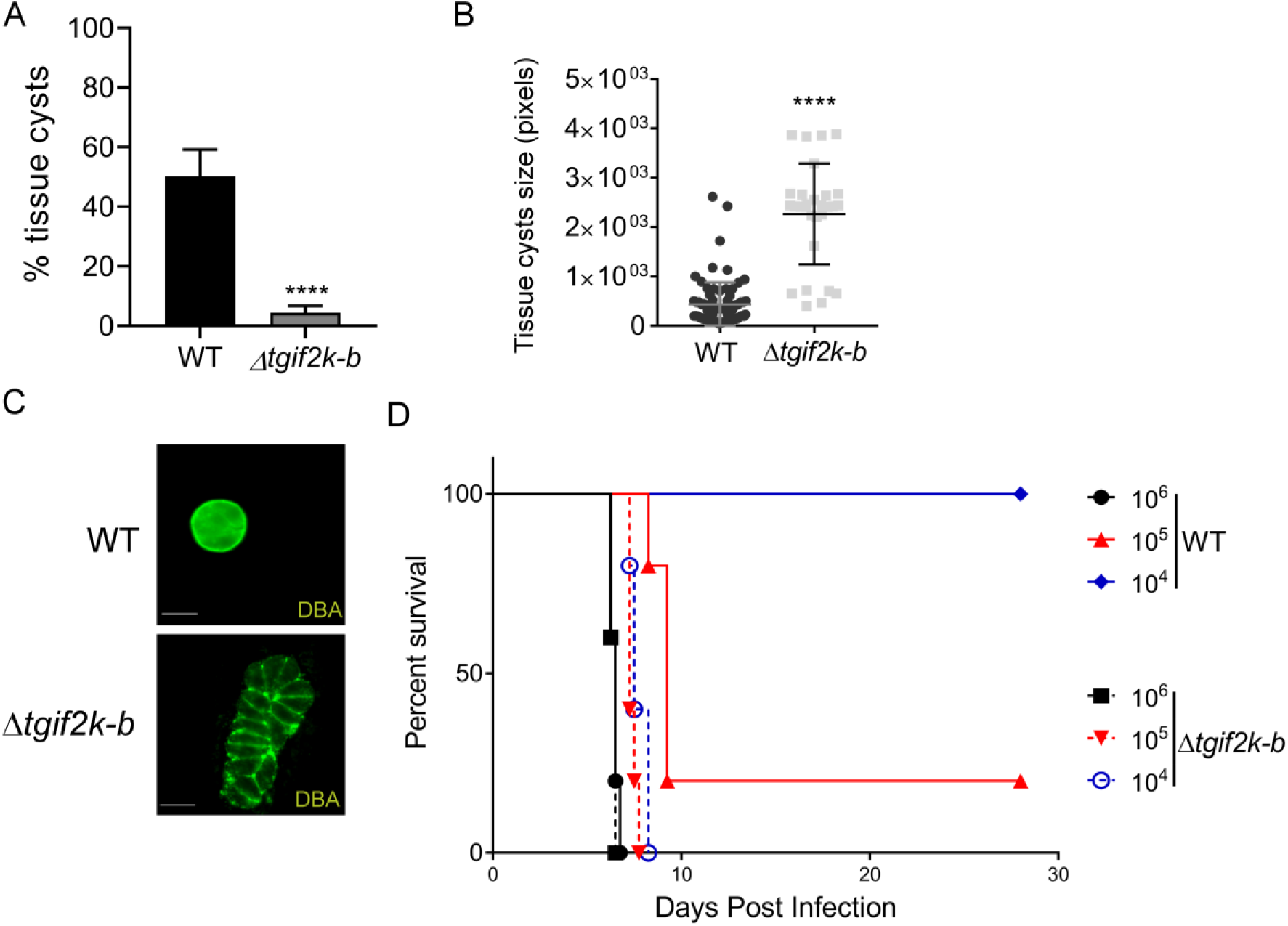
Effects of TgIF2K-B on tissue cyst formation. **(A)** WT or Δ*tgif2k-b* parasites were prompted to spontaneous differentiate in HFF cells. After 5 days, rhodamine-conjugated *Dolichos biflorus* agglutinin (DBA) lectin was used to visualize and count tissue cysts (±SD, n=3). ***p<0.005. **(B)** The area of 100 tissue cysts (in pixels) was measured using ImageJ software (±SD, n=3). ****p<0.001. **(C)** Tissue cysts formed by WT or Δ*tgif2k-b* parasites after 5 days of alkaline stress were visualized by IFA with DBA lectin (green). Scale bar = 10 μm. **(D)** Female BALB/c mice (5 per group) were infected by intraperitoneal injection with 10^4^, 10^5^, and 10^6^ of WT or Δ*tgif2k-b* tachyzoites as indicated. Infected mice were monitored at least 3 times per day and percent survival recorded.

The RNA-seq analysis reveals mechanisms that likely contribute to the growth phenotype observed in Δ*tgif2k-b* parasites, which display bias towards the lytic cycle over differentiation. A transcription factor that represses bradyzoite gene expression, *AP2IX-9* (22), is upregulated in Δ*tgif2k-b* parasites. A master regulator of bradyzoite formation, *TgBFD1* (23), was downregulated in Δ*tgif2k-b* parasites. Bradyzoite-specific genes like LDH2 were also decreased in Δ*tgif2k-b* parasites relative to WT parasites. In combination with the observed higher replication rates of Δ*tgif2k-b* parasites, these results suggest that loss of TgIF2K-B dysregulates the transcriptional program essential for trigger parasite differentiation.

To address the importance of TgIF2K-B *in vivo*, we infected BALB/c mice with increasing doses of WT or Δ*tgif2k-b* tachyzoites. The lack of TgIF2K-B had a profound effect on pathogenesis in the mouse model: BALB/c mice tolerate a dose of 10^4^ WT parasites, but none of the mice were able to survive the equivalent dose of Δ*tgif2k-b* tachyzoites **(Fig. 7D)**. These results mirror the augmented replication rate of Δ*tgif2k-b* tachyzoites seen *in vitro* and further underscore the relevance of TgIF2K-B as a sensor of *in vivo* signals that trigger conversion to latent forms.

## Discussion

The ability of *Toxoplasma* to persist in its host as latent tissue cysts is a key feature that has greatly enhanced the transmission of the parasite. Tissue cysts remain infectious and allow the parasite to infect new hosts via predation. Conversion of tachyzoites to bradyzoites can occur spontaneously in certain cell backgrounds (24) or following exposure to a wide variety of cellular stresses (3). We previously linked the phosphorylation of parasite eIF2α with stress-induced bradyzoite development and latency stages in *Toxoplasma* and fellow apicomplexan parasite *Plasmodium* (9). Four eIF2α kinases have been identified in *Toxoplasma* that respond to ER and nutrient stresses, but the function of TgIF2K-B, which has no clear counterpart in other eukaryotes, had yet to be resolved. Here, we report the discovery that TgIF2K-B contributes to the management of the parasite’s response to oxidative stress.

ROS are a natural consequence of cellular metabolism and have been reported to regulate cellular growth rate. Cells must have systems in place to sense ROS and adapt accordingly. As arsenicals induce ROS that generate oxidative stress, we examined the response of *Toxoplasma* to sodium arsenite. Exposure of the parasites to arsenite induced translational control as evidenced by TgIF2α phosphorylation and diminished protein synthesis **(Fig. 2)**. Using a CRISPR/Cas9 approach, we successfully generated a genetic knockout of TgIF2K-B, which failed to phosphorylate TgIF2α in response to oxidative stress. The ability to initiate translational control in response to ER and nutritive stresses remained unaffected in Δ*tgif2k-b* parasites.

Δ*tgif2k-b* parasites exhibited a faster replication rate relative to WT. We performed an RNA-seq under normal culture conditions and found abnormal transcriptome changes in Δ*tgif2k-b* parasites that featured gene networks involved in protein synthesis and cellular metabolism, consistent with the enhanced parasite replication that was also observed **(Fig. 1E and 3)**. We postulated that the accelerated growth may stem from increased levels of ROS, which have been reported to serve as secondary messengers that influence replication rate in other species (17). Notably, Δ*tgif2k-b* parasites showed significant increases in ROS in the absence of added stress **(Fig. 4A)**. The precise association between ROS levels and proliferation rate is an unresolved question for future investigation.

Abnormal alterations in gene expression occurred between WT and Δ*tgif2k-b* parasites in response to arsenite treatment as well. RNA-seq analysis revealed that loss of TgIF2K-B resulted in dysregulation of oxidation-reduction processes upon arsenite exposure **(Fig. 5)**. The data suggest that without TgIF2K-B, parasites fail to sense oxidative stress and consequently do not slow growth to adapt to this response. Such a phenotype would not be expected to be viable over time; indeed, Δ*tgif2k-b* parasites have been observed to revert to normal growth rates after prolonged culture through an unresolved mechanism (data not shown). Together, these results indicate that TgIF2K-B is central for the management of oxidative stress in *Toxoplasma*, a process that is linked to parasite replication.

It is curious that with loss of translational control and sharply altered mRNA expression, Δ*tgif2k-b* parasites showed increased replication in infected HFF cells **(Fig. 1E)** and in activated macrophages **(Fig. 6)**. There was also enhanced morbidity in mice infected with Δ*tgif2k-b* parasites compared with WT, consistent with the faster replication of the former **(Fig. 7D)**. Our RNA-seq provides mechanistic insights into why tachyzoite replication is enhanced and cyst formation diminished in Δ*tgif2k-b* parasites. For example, *AP2IX-9* is a transcriptional repressor that restricts development of the tissue cyst; its increased expression in Δ*tgif2k-b* parasites would inhibit the expression of bradyzoite mRNAs. Similarly, *TgBFD1* is a master regulator that drives bradyzoite differentiation; its decreased expression in Δ*tgif2k-b* parasites would be expected to impair tissue cyst formation. Consistent with this observation, the loss of TgIF2K-B significantly lowered the frequency of tissue cyst conversion *in vitro;* moreover, the few cysts that did form were larger in size and contained more parasites **(Fig. 7)**.

These results indicate that TgIF2K-B contributes to growth plasticity that enables *Toxoplasma* to appropriately lower the replication rate in response to oxidative stress. To achieve stress adaption, there is an inverse correlation in unicellular eukaryotes between growth rate and stress survival (5). Diminished growth rates during stress are critical for appropriate modulation of gene expression and redistribution of resources to best mediate stress tolerance. In the case of *Toxoplasma*, a critical mode of stress adaptation involves differentiation into bradyzoite cysts, a clinically relevant process that compromised in Δ*tgif2k-b* parasites.

The phenotypes associated with loss of TgIF2K-B are representative of antagonistic pleiotrophy (25), where the gene regulates multiple traits. Enhanced replication is a beneficial trait associated with Δ*tgif2k-b* parasites, but it comes at the expense of the parasite being able to modulate proliferation when faced with signals to transition into latent stages. Therefore, TgIF2K-B functions as a critical sensor to balance parasite replication and latency; parasites lacking TgIF2K-B would likely have reduced transmission through the predation route due to their compromised ability to form infectious tissue cysts.

## Materials and Methods

### Parasite culture

ME49 tachyzoites were cultivated in human foreskin fibroblast (HFF, ATCC) monolayers in Dulbecco modified Eagle medium (DMEM) supplemented with 5% heat-inactivated fetal bovine serum (FBS) (Gibco/Invitrogen), 100 U/ml penicillin, and 100 μg/ml streptomycin. HFF cells were cultivated in DMEM supplemented with 10% FBS. The cultures were maintained in a humidified incubator at 37°C with 5% CO_2_.

### Disruption of TgIF2K-B genomic locus

The Δ*tgif2k-b* parasites were generated by disrupting the genomic locus using CRISPR/Cas9 and an associated guide targeting the sequence ACCTCTCGCTCTGCGTTCCCT to allow integration of a minigene encoding a modified dihydrofolate reductase (DHFR) that confers pyrimethamine resistance (15, 16). The pSAG1:U6-Cas9:sgRNA-TgIF2K-B vector was generated using Q5 site-directed mutagenesis (New England Biolabs) (26) and the listed forward and reverse primers **(Table S1)**. After transfection, the parasites were selected using 1 μM pyrimethamine for three passages before cloning by limited dilution in 96 well-plates. Clones were selected and assayed by PCR using primer pairs indicated in Figure 1A.

### Immunoblot for TgIF2K-B

Parasites were purified from HFF cells using a 3 micron filter and lysates were made by suspension of the parasites pellets in a PBS solution supplemented with 0.01% Triton X-100. Proteins in parasite lysates were resolved NuPAGE using 4-12% gradient bis-Tris polyacrylamide gels (Thermo Fisher Scientific). Separated proteins were then transferred from the gels to a nitrocellulose membrane. Immunoblotting was performed using antibody specific to TgIF2K-B (8) diluted 1:10 in blocking solution (Tris-buffered saline (TBS) pH 7.4 with Tween 20 and 2.5% BSA). The secondary antibody, goat anti-Rabbit IgG (H+L) Secondary Antibody, HRP, was used at dilution of 1:1,000. After washing, the membrane was incubated with Pierce enhanced chemiluminescence (ECL) immunoblot substrate to visualize the proteins levels using the FluorChem E system (Protein Simple).

### Measurements of mRNA levels

RNA was isolated from parasites using TRIzol LS (Invitrogen) and cDNA was generated using Omniscript (Qiagen). RT-qPCR was carried out using primers specific (Table S1) with SYBR Green Real-Time PCR Master Mixes (Invitrogen) and the StepOnePlus Real System (Applied Biosystems). Relative levels of transcripts were calculated with the ΔΔ*Ct* method using the *GAPDH* gene as an internal control. Each experiment was performed three times, each with three technical replicates.

### Measurements of TgIF2α phosphorylation

Tachyzoites were purified from infected HFFs by syringe passage and filtration as described in (27) and then incubated in DMEM supplemented with 5% FBS in the presence of a stress agent or vehicle: 1 μM thapsigargin (TG, Sigma-Aldrich), 50 nM halofuginone (HF, Sigma-Aldrich), or 5 μM sodium arsenite (Ars, Sigma-Aldrich), for 2 h at 37°C. For some reactions, 20 μM N-acetyl cysteine (Nac, Sigma-Aldrich) was added. Parasites were lysed in PBS containing 0.01% Triton X-100, supplemented with the protease inhibitor cocktail cOmplete and an EDTA-free protease inhibitor cocktail (Sigma-Aldrich). Total protein levels were quantified using Pierce BCA Protein Assay Kit (Thermo-Fisher). Proteins were separated by NuPAGE 4-12% bis-Tris gels (Thermo Fisher Scientific) and transferred to a nitrocellulose membrane for immunoblot analyses using antibodies recognizing total TgIF2α or phosphorylated TgIF2α (1:1000) (27). The secondary antibodies, goat anti-Rabbit IgG (H+L) Secondary Antibody, HRP, were used at 1:5,000 dilution. After washing, the membranes were incubated with Pierce ECL substrate to visualize the proteins.

### Parasite replication assays

Tachyzoites were allowed to invade HFF monolayers for 2 h. Infected cultures were then washed and incubated with fresh culture medium. At 36 h post-infection, infected HFF monolayers were fixed with 2.5% paraformaldehyde for 20 min, then blocked with PBS supplemented with 2% BSA. Cells were incubated with anti-SAG1 (Invitrogen) in blocking buffer containing 0.2 % Triton X-100 for 1 h. Next, goat anti-mouse Alexa-Fluor 488 (Invitrogen) was applied for 1 h, followed by ProLong Gold Antifade Mountant with DAPI to visualize parasites. The number of parasites in 250 randomly selected vacuoles was counted for each sample.

J774.1 macrophages (ATCC) were incubated with 10 ng/ml of LPS for 24 h before being infected with tachyzoites for 2 h at an MOI of 3 (28). Following 4 days of infection, cells were harvested and the numbers of parasites were quantified using the PCR-based B1 assay (21). To test parasite fitness in HFFs after cycling through macrophages, and infected macrophages were harvested by scrape and syringe lysis at 2 days post-infection. The same volume of media containing tachyzoites was then used to infect an HFF monolayer for 2 h, which was then washed with DMEM twice. Six days post-infection, the degree of host cell lysis was determined by crystal violet staining as described previously (29).

### Puromycin incorporation

To determine the levels of protein synthesis in WT and Δ*tgif2k-b* parasites, extracellular tachyzoites were subjected to 5 μM arsenite for 2 h. Parasites were then incubated with 10 μg/mL puromycin (Sigma) for 15 min and parasites were lysate in PBS+ 0.01% Triton X-100. Total protein synthesis was measured by immunoblot using the anti-puromycin antibody at 1:500 (EMD Millipore). Protein synthesis was quantified by densitometry using ImageJ and normalized by immunoblot using antibody specific to steady-state levels of eIF2α protein (13).

### RNA preparation and sequencing

Tachyzoites were allowed to invade HFF host cell monolayers for 2 h, then infected monolayers were washed and replaced with fresh medium. At 36 hours post-infection, intracellular tachyzoites were harvested from HFFs by syringe passage and filtering. Then parasites were washed in DMEM and centrifuge 300g for 15 min. Purified extracellular WT and Δ*tgif2k-b* tachyzoites were treated with vehicle or 5 μM sodium arsenite for 2 h at 37°C and total RNA was extracted using RNeasy (Qiagen). The RNA concentration for each sample was measured using Nanodrop One (Thermo Scientific). Library preparation and sequencing was performed by GENEWIZ (South Plainfield, NJ). The quality of the raw sequencing reads obtained from GENEWIZ was checked using FastQC (30) and the low-quality reads were trimmed using TrimGalore (31). Trimmed reads were aligned to the reference *T. gondii* ME49 genome (ToxoDB.org) using hierarchical indexing for spliced alignment of transcripts (HISAT2) (32). HISAT2 analysis was performed using the high-throughput computing cluster, Karst, which is available at Indiana University. The number of reads aligning to the exons of different genes were counted using featureCounts (33). Lastly, differential expression analysis was carried out using DESeq2 (34). Volcano plots were created using R (35), and genes with log2fold change of ≥ ±1 and p-adjusted value of ≤ 0.05 were considered significant for the further network analyses. Gene Ontology enrichment analyses were performed using the collated genes that were significantly upregulated or downregulated, with a focus on molecular and biological cellular processes. RNA-seq datasets from this study are available in the NCBI GEO database (accession # GSE158231).

### Bradyzoite differentiation assays

ME49 tachyzoites were allowed to infect HFF monolayers for 2 h. After washing and replacing DMEM 5% FBS. The cells were cultured at 37°C for 5 days. To visualize tissue cyst walls, infected cells were fixed with 2.5% paraformaldehyde and stained with rhodamine-conjugated with *Dolichos biflorus* agglutinin (Vector laboratories). Tissue cysts were counted in 250 random infected cells. Also, the tissue cyst area was measured using ImageJ by drawing a region of interesting (ROI).

### Reactive oxygen species

ROS were measured using the OxiSelect *In Vitro* ROS/RNS Assay Kit (Cell Biolabs, Inc.) per the manufacturer’s instructions. Briefly, lysates made from 10^7^ parasites were added to a 96-well plate, followed by 50 μl of Catalyst reagent of the kit. Reactions were incubated for 5 minutes at room temperature before adding 2’,7’-Dichlorofluorescin (DCFH) solution into each well for 1 h more incubation. Fluorescence was measured at 480 nm/530 nm Ex/Em in a Synergy H1 microplate reader (BioTek). All samples were assayed in triplicate.

### Analysis of infection in mice

Five to 6-week-old female BALB/c mice were purchased from Jackson Laboratory (Bar Harbor, ME) and allowed to acclimate in our facility for 1 week. Following acclimation, the mice were randomized on the basis of weight into six groups (n = 5). WT or Δ*tgif2k-b* tachyzoites were prepared in sterile PBS and intraperitoneally injected into the mice, which were then monitored at least 3 times a day. The mice used in this study were housed in American Association for Accreditation of Laboratory Animal Care (AAALAC)-approved facilities at the IUSM Laboratory Animal Research Center (LARC). The Institutional Animal Care and Use Committee (IACUC) at Indiana University School of Medicine has approved the use of all animals and procedures (IACUC protocol number 11376).

### Quantification and statistical analysis

Quantitative data are presented as the mean and standard deviation from biological triplicates. Statistical significance was determined using One-way ANOVA with Tukey’s post hoc test or multiple t-test two tailed in Prism (version 7) software (GraphPad Software, Inc., La Jolla, CA). The number of biological replicates (n) and p values are indicated in the legend of each figure. For immunoblot analyses, the reported images are representative of at least three independent experiments.

## Acknowledgments

This research was supported by a research grant from National Institutes of Health (AI124723 to W.J.S. and R.C.W.). J.M. was supported by PHS grant T32 AI060519 and the Joseph and Lucille Madri Family Scholarship. R.C.W. has received grant support from Eli Lilly and Company and is a scientific advisor to HiberCell. Other authors declare no conflicts of interest. The authors would like to thank the Biology of Intracellular Pathogens Group at IUSM and other members of the Wek laboratory for helpful discussions and suggestions.

## Supplemental Figure Legends

**Supplemental Figure 1.**
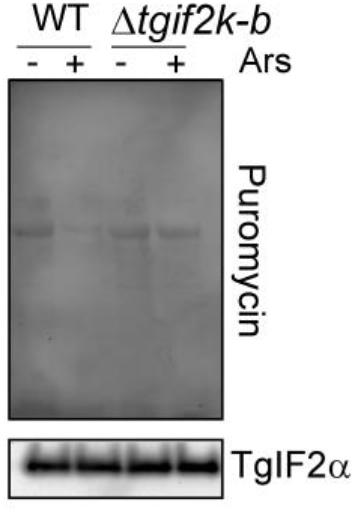
Loss of TgIF2K-B increases protein synthesis. WT and *Δtgif2k-b* tachyzoites were treated with 5 μM sodium arsenite (Ars) for 2 hours. Global translation was measured by incubating cells with puromycin for 15 min, followed by lysate preparation and immunoblot analyses with puromycin-specific antibodies.

**Supplemental Figure 2.**
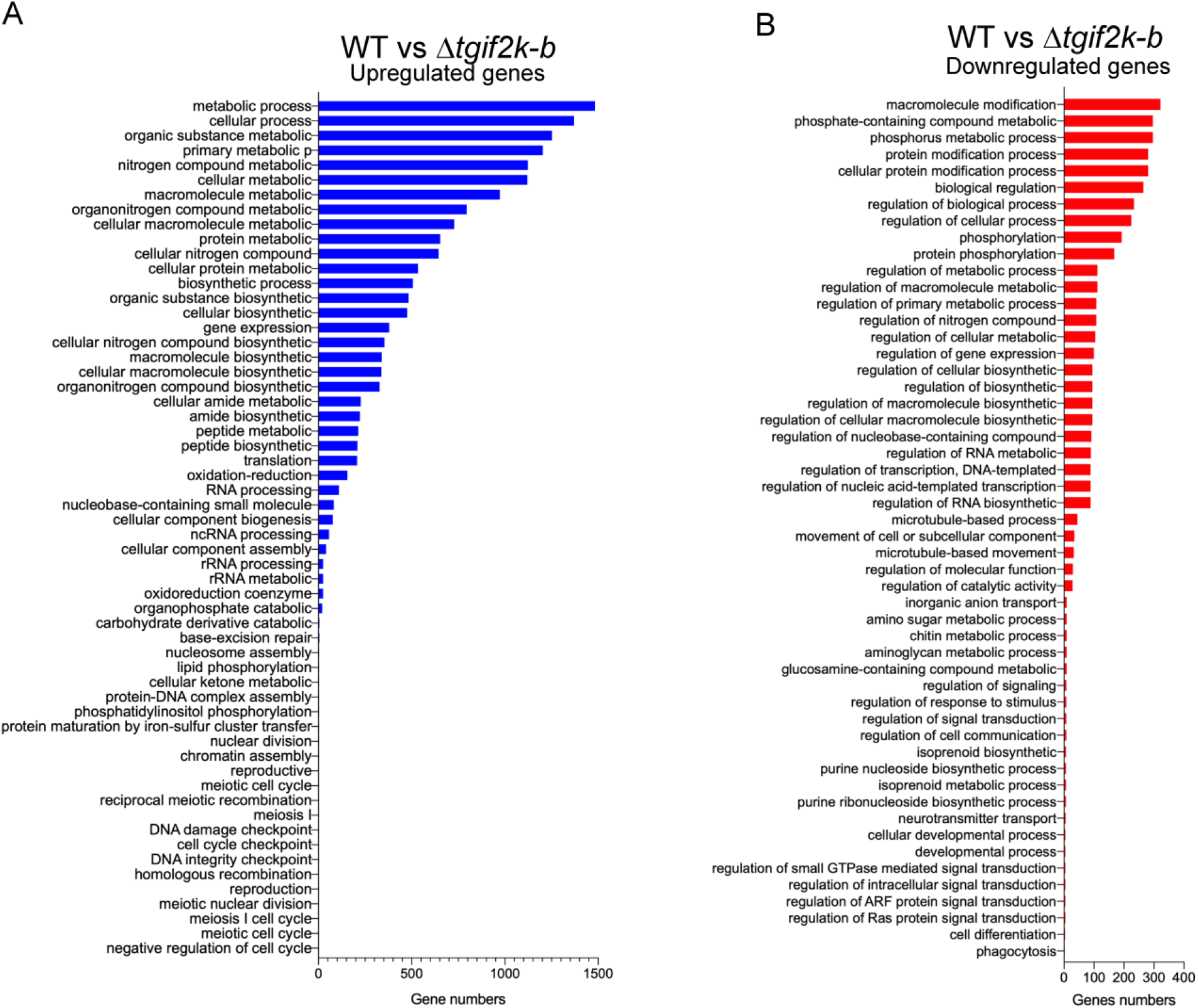
Gene ontology enrichment analysis for biological processes. Gene ontology enrichment analysis for biological processes (using ToxoDB) of **(A)** upregulated genes and **(B)** downregulated genes in *Δtgif2k-b* compared to WT parasites. The length of the bars indicates the number of genes identified for the indicated processes.

**Supplemental Figure 3.**
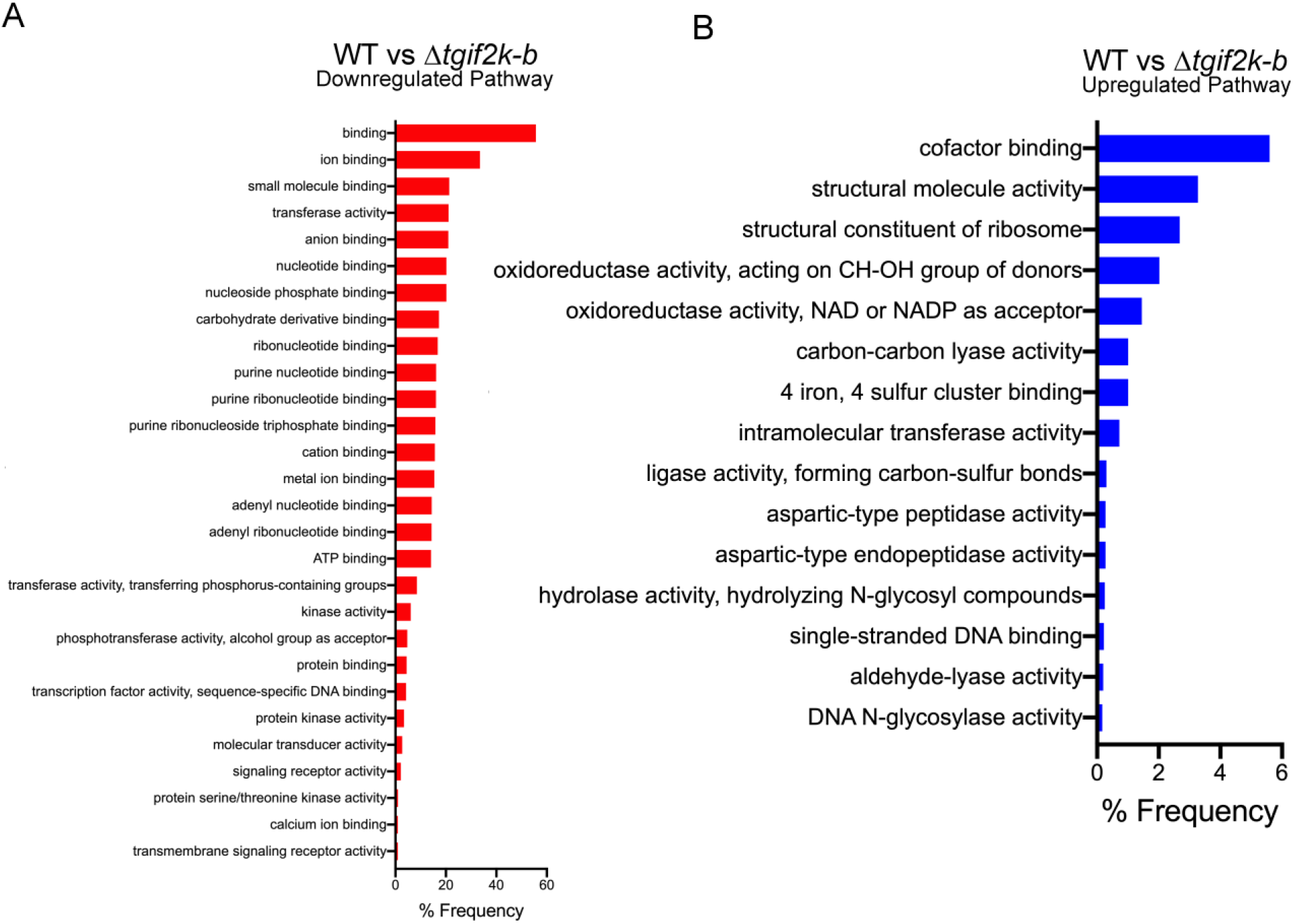
Gene ontology enrichment analysis for molecular function. Gene ontology enrichment analysis for molecular function (using ToxoDB) of (A) upregulated genes and (B) downregulated genes in *Δtgif2k-b* compared to WT parasites. The length of the bars indicates the number of genes identified for the indicated processes.

## Supplemental Tables

**Table S1.** Primer sequences.

**Table S2.** RNA-seq analysis of *Δtgif2k-b* parasites compared to WT.

**Table S3.** RNA-seq analysis of WT parasites (untreated versus arsenite-treated).

**Table S4.** RNA-seq analysis of *Δtgif2k-b* parasites (untreated versus arsenite-treated).

**Table S5.** Metabolic Pathway Enrichment of downregulated genes in WT (untreated versus arsenite-treated).

